# Microgeography of staphyloccoci in human tissue explains antibiotic failure

**DOI:** 10.1101/2025.09.12.675782

**Authors:** Vishwachi Tripathi, Benedict Morin, Minia Antelo Varela, Aline Amani Loriol, Kristiina Kurg, Sandro Jakonia, Asvin Ehsan, Sébastien Herbert, Alexia Ferrand, Loïc Sauteur, Florian C. Marro, Noemi Henselmann, Jasmin Künnecke, Fabienne Estermann, Kerstin Strenger, Aya Iizuka, Anne-Kathrin Woischnig, Beatrice Claudi, Sandra Söderholm, Peter M. Keller, Daniel Baumhoer, Thomas Bock, Krittapas Jantarung, Pablo Riviera Fuentes, Sylvain Meylan, Mario Morgenstern, Martin Clauss, Nina Khanna, Richard Kuehl, Dirk Bumann

## Abstract

Bacterial infections remain a major health threat, yet pathogen biology in human tissues is poorly understood. Using AI-guided imaging, we mapped ∼15,500 *Staphylococcus aureus* cells in biopsies from 33 patients undergoing surgery for musculoskeletal infections. Despite substantial interindividual variability, consistent patterns emerged. Most bacteria resided within non-classical monocytes/macrophages, challenging models of primarily extracellular pathogenesis. Both intra- and extracellular bacteria were predominantly isolated single cells or doublets with low rRNA content, suggesting limited replication. Complementary proteomics implicated inflammation-associated hypoxia and host glucose-to-lactate metabolism as growth constraints. Preoperative antibiotic therapy failed to clear bacteria across microenvironments and cluster sizes, challenging assumptions that antibiotic tolerance is confined to intracellular niches or biofilms and underscoring the clinical need for debridement. In vitro models replicating diverse tissues conditions impaired antibiotic activity, indicating multifactorial resilience. Together, these findings redefine *S. aureus* infection biology in musculoskeletal infections and establish a framework for mechanism-based prevention and therapy.

## Main

The rise of antimicrobial resistance limits treatment options for bacterial infections, and effective vaccines remain unavailable for several major pathogens. Global travel and increasingly complex medical procedures accelerate pathogen spread, highlighting the need for novel anti-infective strategies ^1,2^. Most research and development relies on artificial laboratory conditions that promote rapid bacterial growth. However, animal models and sparse human data indicate that, within host tissues, bacteria can adopt markedly different phenotypes, including slow growth, biofilms, heterogeneous subpopulations, and intracellular niches ^3–5^. Such adaptations would affect antibiotic activity and immune effector mechanisms, requiring tailored control strategies. However, their relevance in human disease remains unclear ^6–8^, because comprehensive, multi-patient, spatial datasets from human tissues are lacking across bacterial pathogens. Critical obstacles include limited access to human biopsies, the need to detect sparse micrometer-sized bacteria in centimeter-scale tissues, high autofluorescence in inflamed tissues, and extensive tissue- and patient-level heterogeneity ^9–11^.

Here, we addressed these challenges in human musculoskeletal infections caused by *Staphylococcus aureus*, a leading global cause of morbidity and mortality ^1,2,12^. These infections frequently require surgical debridement ^5,12–16^, because antimicrobial chemotherapy fails despite *S. aureus* susceptibility and sufficient drug penetration at the infection site ^17^. Clinical vaccine trials to prevent such infections have failed to show protective efficacy. Multiple models could explain these prevention and treatment failures ^5,12,18–21^, but direct evidence from infected human tissues is sparse.

### Feedback microscopy enables efficient *S. aureus* localization in human tissues

We imaged intraoperative tissue samples from a total of 33 individuals undergoing surgical debridement for monomicrobial *S. aureus* musculoskeletal infections. Nineteen individuals were included in the initial analysis (Fig. 1a,b) and 14 subsequently added for validation with a different sample processing and imaging pipeline (Extended Data Table 1). We included patients with a broad spectrum of acute or chronic infections, including native joint arthritis, fracture-related infections, osteomyelitis without implant, soft-tissue infections, and prosthetic joint infections. Some patients received preoperative antibiotics. Tissue specimens encompassed muscle, connective tissue, fibrinopurulent material, and granulation or scar tissue. Whole-genome sequencing of 21 *S. aureus* strains isolated from parallel biopsies revealed genetic diversity covering major global lineages ^22^ (clonal complexes 1, 5, 8, 22, 30, 45, 97, and additional serotypes 7, 101, 121). Thus, patients, infection types, antibiotic exposure, tissue context, and bacterial strains were heterogeneous (Fig. 1b)^23^. Our goal was to determine whether common aspects of *S. aureus* infection biology existed across this diversity.

**Figure 1:**
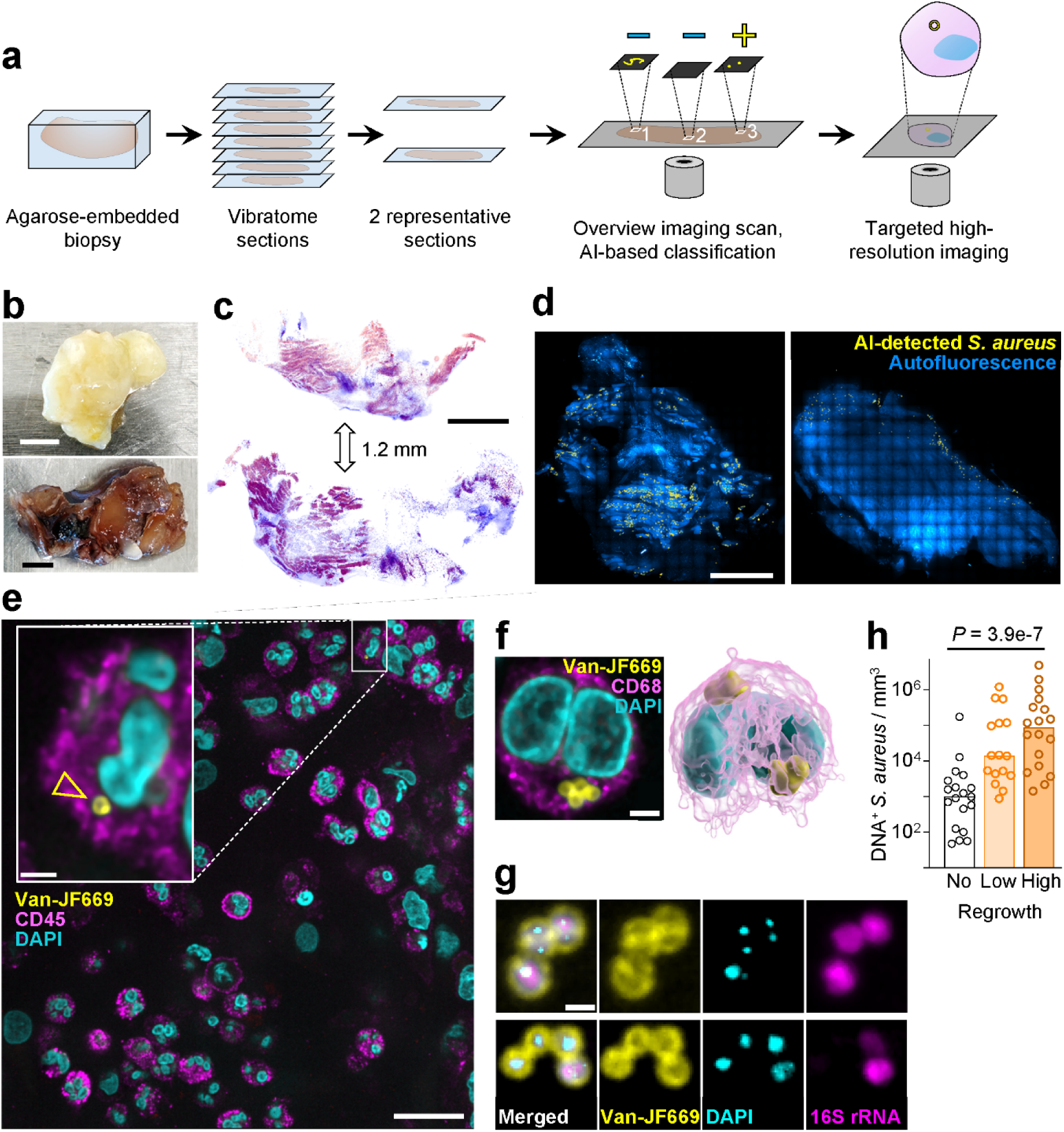
Visualizing *S. aureus* in human tissues. **(a)** Schematic overview of imaging pipeline. Fixed patient biopsies are embedded in agarose and serially sectioned with a vibratome. Two representative sections are selected for overview microscopy scans. AI-segmentation identifies background signals (-) and possible *S. aureus* (+) for targeted imaging of the bacteria at higher resolution. **(b)** Biopsies. Scale bar, 5 mm. **(c)** Two sections from the same biopsy spaced 1.2 mm apart. The sections were stained with a histology-emulating fluorescence method. Scale bar, 1 mm. **(d)** Positions of possible *S. aureus* objects in two overview scans from different biopsies. Scale bar, 1 mm. **(e)** Context of a *S. aureus-*containing immune cell (magnified in the inset). Scale bars, 20 μm and 2 μm (inset). **(f)** *S. aureus-*containing immune cell and 3D reconstruction from a confocal stack. Scale bar, 2 μm. **(g)** Visualization of DNA stained with DAPI and rRNA stained with a fluorescent probe to 16S rRNA in *S. aureus* cells identified with fluorescent vancomycin (Van-JF669). Scale bar, 1μm. **(h)** Relationship between semi-quantitative ex vivo bacterial growth results (Med/Hi, medium/high) and density of DNA-containing *S. aureus* based on imaging for two halves of the same biopsies. One-way ANOVA of log-transformed data with post-test for linear trend.

Imaging 5 μm sections of formalin-fixed paraffin-embedded (FFPE) biopsies with commercially available vancomycin-BODIPY ^24,25^ revealed only few *S. aureus* and high tissue autofluorescence ^26^ (Extended Data Fig. 1a). Autofluorescence suppression (Extended Data Fig. 1b) ^27^, replacement of FFPE sections with 60 µm agarose-embedded vibratome sections (Fig. 1a,c; Extended Data Fig. 1e), and staining with vancomycin-Janelia Fluor 669 (Van-JF669) ^28^ enabled robust detection of numerous *S. aureus* with excellent signal-to-background ratio, except in regions with fibers emitting broad-spectrum autofluorescence (Fig. 1d-f; Extended Data Fig. 1c, d).

Imaging a single tissue section at a 3D resolution sufficient for bacterial detection required >24h, even with a speed-optimized spinning-disk confocal microscope. The resulting datasets exceeded 1 Terabyte and often contained extensive tissue regions without bacteria (Fig. 1d; Extended Data Fig. 1f). We reduced imaging time and data volume by targeting high-resolution acquisition only to relevant areas using feedback (“smart”) microscopy ^29–32^ (Fig. 1a; Extended Fig. 1c). An initial moderate-resolution overview scan was followed by rapid AI-based identification of bacteria and immediate targeted re-imaging of corresponding regions at high resolution. AI-based detection achieved >95% sensitivity for bacteria and avoided visualization bias toward large clusters (as in direct manual inspection).

Multiple sections from the same biopsy frequently revealed heterogeneous tissue architecture and patchy infection (Fig. 1c, d; Extended Data Fig. 1e, f). To account for intra-sample heterogeneity, we imaged sections at every 900 µm in z-direction (orthogonal to the sectioning plane) using a virtual hematoxylin and eosin staining compatible with thick sections ^33^ (Fig. 1c; Extended Data Fig. 1e) and selected the two most diverse sections per biopsy.

The majority of Van-JF699-labelled *S. aureus* cells contained 4’,6-diamidino-2-phenylindole (DAPI)-stained DNA (median 69%, IQR 35–93%) (Fig. 1g). *S. aureus* lacking DNA likely represented dead bacteria that had lost cytosolic contents ^34^, while Van-JF669-stained peptidoglycan ^35^ remained structurally intact ^34^ until cleared by host enzymes ^36^. The density of DNA^+^ *S. aureus* cells in a section correlated with ex vivo growth of *S. aureus* from an unfixed part of the same biopsy (Fig. 1h), supporting DNA as a viability marker. To assess ribosome content, we employed fluorescence in situ hybridization (FISH) targeting *S. aureus* 16S rRNA ^37–39^ (Fig. 1g). To determine bacterial microenvironments, we stained host markers including CD14, CD16, CD45, CD68, calprotectin, vimentin, and seven cytosolic proteins (Fig. 1e, f; Extended Data Table 2). We report only results for samples with ≥50 detected bacteria to ensure representative data.

Adapting our imaging pipeline to FFPE sections enabled automated detection of *S. aureus*, DNA, and ribosomal content also in standard histopathology samples (Extended Data Fig. 1g). We included a second cohort of 14 patients using this alternative sample processing and imaging pipeline, to validate our findings about intra-/extracellular states as we as bacterial density and aggregation.

### Dominance of intracellular and isolated *S. aureus* cells

Imaging of biopsies from 33 patients (primary and validation cohorts) revealed 15,448 *S. aureus* cells. Bacterial density varied greatly between and within biopsies (Fig. 1d, e, h; Ext Data Fig. 1f). All patients harbored both extra- and intracellular *S. aureus*, with intracellular bacteria predominant in most individuals from both cohorts (median 73%, IQR 60–84%; Fig. 1f; Fig. 2a, b). Intra- and extracellular bacteria had similar DNA content, suggesting comparable viability (Fig. 2b). Thus, intracellular niches emerged as a major part of *S. aureus* infection biology. Most infected cells contained only one or two bacteria (Ext Data Fig. 1h). In all analyzed patients, >95% of infected cells were CD45^+^ immune cells. In most patients, these cells were primarily CD16^+^ CD68^+^ (median 70.6%, IQR 50.3-81.5%; Fig. 2c, d), and ∼60% of these were also positive for CD14 and calprotectin, consistent with non-classical activated monocytes differentiating into macrophages ^40,41^. Uninfected cells of this type were common in several biopsies (Ext Data Fig. 1i), suggesting no specific targeting by *S. aureus*. Other infected cells included CD16^+^ CD68^−^ granulocytes (median 11%, IQR 3–18.8%) and CD16^−^ CD68^+^ macrophages (median 11.3%, IQR 2.9–18.1%), indicating that most infected host cells were professional phagocytes. Infected cells displayed intact nuclei, compromised nuclear membranes, or cell lysis with extracellular release of DNA and calprotectin entrapping *S. aureus* (Fig. 2e, f, g). This suggested cell death with expulsion of extracellular traps (ETosis) as a mechanism for release of intracellular *S. aureus* ^42–45^.

**Figure 2:**
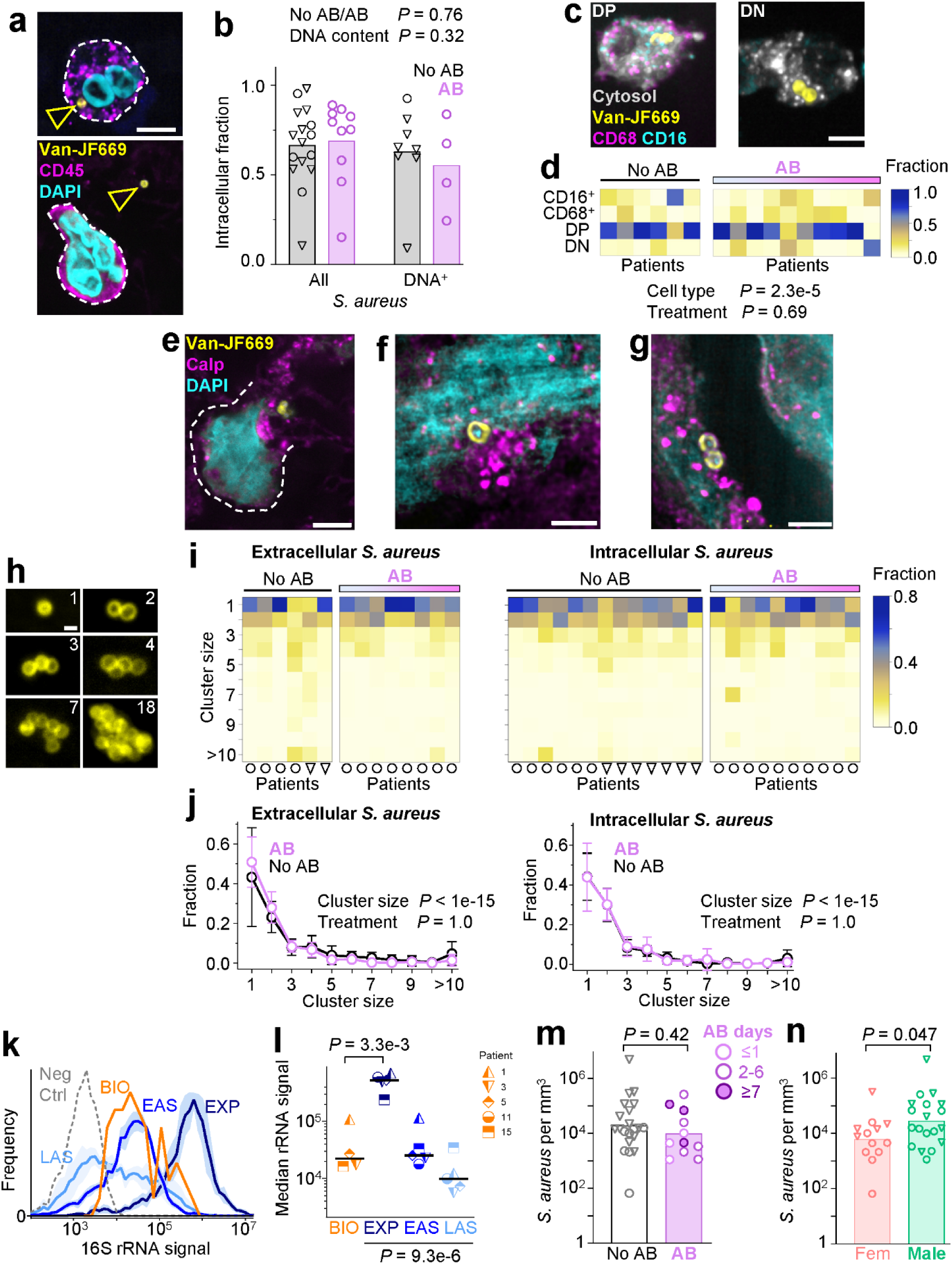
*S. aureus* phenotypes in human tissues. **(a)** Examples of an intra- (top) and an extracellular (bottom) *S. aureus* (arrowheads) in or close to immune cells. Scale bar, 5 μm. **(b)** Intracellular fraction of all *S. aureus* or DNA-containing *S. aureus* in patients treated or not with antibiotics (AB). Each circle represents one patient (circles, cohort 1; triangles, cohort 2); the bars represent the means. Two-way ANOVA for impact of antibiotics and DNA content. We only included data with >50 bacteria per sample (as in all other figures). **(c)** Examples of an infected CD16^+^ CD68^+^ double-positive (DP, left) and an infected double-negative (DN, right) cell. Scale bar, 5 μm. **(d)** Distribution of infected cell types. Each column represents one patient. Patients receiving antibiotics are sorted according to length of prior treatment. Two-way ANOVA for impact of cell type and antibiotic treatment. Example of a lysing infected cell (Calp, calprotectin). Scale bar, 5 μm. **(e)-(g)** Examples of extracellular *S. aureus* entangled in DNA and calprotectin (Calp), **e** shows a lysing cell with the cell perimeter marked by the dashed line. Scale bars, e, 5 μm; f, g, 2 μm. **(h)** Examples of objects with different *S. aureus* numbers. Scale bar, 1 μm. **(i)** Distribution of *S. aureus* cluster sizes outside (left) and inside (right) of host cells. Each column represents one patient (circles, cohort 1; triangles, cohort 2). Patients receiving antibiotics (AB) are sorted according to length of prior treatment. **(j)** Comparison of cluster size distribution in patients treated or not with antibiotics (AB). Means and SD of 6 to 13 patients are shown. Two-way ANOVA. **(k)** Distribution of 16S rRNA signals of *S. aureus* in biopsies (BIO, pooled from 5 patients) and in vitro cultures at exponential (EXP, 2h after inoculation), early stationary (EAS, 16h), or late stationary (LAS, 48h) growth phases (strain from patient 3). Signals for a control probe with inverted sequence are also shown (Neg Ctrl). **(l)** Median 16S rRNA signals of *S. aureus* in biopsies (BIO) and in vitro cultures of the same isolates at exponential (EXP), early stationary (EAS), or late stationary (LAS) growth phases. Each symbol represents signals detected in one patient or the geometric mean of signals observed in three independent in vitro experiments. Comparison of biopsies with exponentially growing cultures, paired t-test of log-transformed data. Comparison of in vitro cultures, one-way ANOVA of log-transformed data with post-test for linear trend. **(m)** Density of *S. aureus* objects (singles and clusters) in biopsies from patients treated with antibiotics (AB) or not. Each symbol represents one patient (circles, cohort 1; triangles, cohort 2); the bars represent geometric means. t-test of log-transformed data. **(n)** Density of *S. aureus* objects (singles and clusters) in biopsies from female (Fem) and male patients. Each symbol represents one patient (circles, cohort 1; triangles, cohort 2); the bars represent geometric means. t-test of log-transformed data.

*S. aureus* growing planktonically in vitro typically forms grape-like clusters (*staphyle* is Greek for grape). Clusters of >10 bacteria were present in our biopsies (Fig. 2h, Extended Data Fig. 1g), as observed previously ^46–54^, but they were rare. Most bacteria were single cells or connected pairs in both cohorts (Fig. 2h, i, j), suggesting limited replication. This was supported by rRNA levels similar to early stationary-phase *S. aureus* in vitro cultures (comparing isolates from the same patients; Fig. 2k, l). There was no detectable link between aggregate size and DNA content (Ext Data Fig. 1k), arguing against improved survival in clusters. It is important to note that we did not investigate bone and implants, which might harbor large aggregates ^46–54^.

Ten of 32 patients received adequate antibiotics for a few hours to 29 days (median 4 days, IQR 1-5 days) before surgery. We detected *S. aureus* in all biopsies from these patients with comparable densities for untreated and treated patients (Fig. 2m), consistent with the limited in-tissue efficacy of antimicrobial chemotherapy despite antimicrobial susceptibility in laboratory tests and efficient tissue penetration ^17^. Prior models predicted superior survival of intracellular *S. aureus* or *S. aureus* in aggregates ^5,55^, which would be expected to shift bacterial populations toward these subsets in treated patients. However, untreated and treated patients displayed comparable intracellular fractions, host cell types, cluster sizes, and DNA-based viability (Fig. 2b,d,i,j,m; Extended Data Fig. 1h,k). Thus, antibiotic survival extended beyond intracellular bacteria and biofilms/aggregates, with even isolated extracellular cells demonstrating resilience. This broad tolerance was consistent with the need for surgical debridement in treatment of musculoskeletal infections.

Comparison of samples from female vs. male patients revealed no significant sex differences except for a slightly lower bacterial density in female patients (Fig. 2n). Limited sex differences were consistent with findings for *S. aureus* bacteremia ^11^.

Taken together, *S. aureus* showed heterogeneous distributions within and between patients ^56,57^. Across this diversity, common patterns emerged, including coexistence of extra- and intracellular bacteria in all patients, a dominant role of non-classical activated monocytes/macrophages as host cells, and a preponderance of isolated bacteria with limited replication and ribosome content across diverse microenvironments. Antimicrobial survival did not seem to be restricted to a single niche, suggesting broad, multifactorial resilience as another common feature.

### Host inflammation-associated metabolism limits *S. aureus* growth

To identify *S. aureus* activities during infection, we analyzed biopsies snap-frozen directly in the operating room from 12 individuals using targeted proteomics for virulence factors, metabolism, and stress responses ^58^ (Fig. 3a; Ext Data Fig. 2a). *S. aureus* in biopsies upregulated toxins leucocidin A (LukA) and γ-hemolysin (HlgC) but down-regulated EsxA, as reported previously ^59–63^. Leukocidin A can induce rapid host cell ETosis ^64^, consistent with our imaging (Fig. 2f, g) ^64^. Enzymes linked to growth and division, PBP2 and Atl ^65,66^, were markedly downregulated, suggesting slow replication, consistent with the dominance of isolated single bacteria/doublets and low rRNA content (Fig. 2i, k, l).

**Figure 3:**
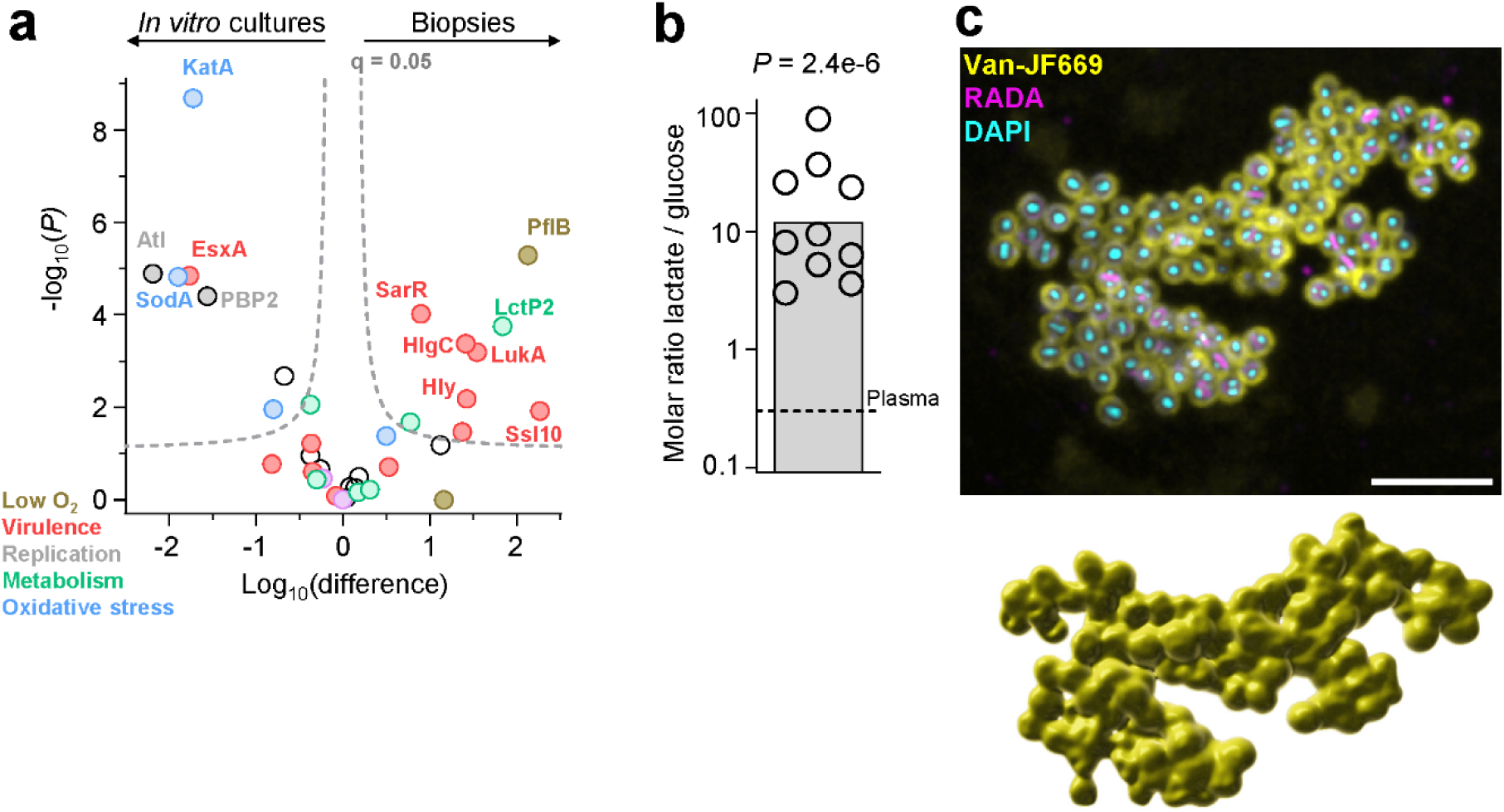
*S. aureus* metabolic restrictions in human tissues. **(a)** *S. aureus* protein abundance in biopsies and in vitro cultures. Each circle represents the geometric mean of 8 exponentially growing in vitro cultures in Mueller-Hinton broth and detected protein in at least 3 biopsies. The dashed line represents a false-discovery rate of 0.05. **(b)** Lactate / glucose ratios in biopsies. Each circle represents one patient; the bar is the mean. The dashed line represents the ratio in plasma. One-sample t-test of log-transformed data against the plasma value. **(c)** *S. aureus* cluster grown in a live biopsy slice on rich medium and normoxia for 16h. A 5 μm thick section of the cluster containing >80 bacteria is shown. The fluorescent D-amino acid RADA was present during the last 30 min of incubation to highlight cell-wall synthesis. The lower panel shows a 3D reconstruction of this cluster. Scale bar, 5 μm.

The low-oxygen marker protein PflB ^67^ was upregulated >100-fold, suggesting limited oxygenation (Fig. 3a; Ext Data Fig. 2a). Such hypoxia often occurs in infected tissues due to oxygen consumption by infiltrating immune cells and thrombosis-driven perfusion deficits ^70–73^. This was consistent with poorly perfused infected tissue in musculoskeletal infections that requires extensive debridement until bleeding from small vessel appears ^68,69^. Under these conditions, immune cells shift from aerobic respiration to glycolysis, converting glucose to lactate ^74^. Lactate indeed accumulated to high levels in the biopsies (Fig. 3b) ^75^, and *S. aureus* upregulated the lactate transporter LctP2 (Fig. 3a) suggesting lactate utilization ^76^. All tested musculoskeletal *S. aureus* isolates could utilize lactate as carbon/energy source with moderate yields in vitro (Extended Data Fig. 2b). In contrast to glucose, lactate is non-fermentable and supports *S. aureus* growth only when electron acceptors such as oxygen are available ^78^. Thus, host glucose-to-lactate conversion together with oxygen depletion likely restricted bacterial growth in infected tissues ^71,79^. Elevated lactate may also enhance the activity of *S. aureus* alpha-hemolysin (Hly, abundant in biopsies, Extended Data Fig. 2a), increasing virulence ^80^.

All tested musculoskeletal isolates *S. aureus* were auxotrophic for various amino acids ^81,82^ (Extended Data Fig. 2c). Arginine, required by all isolates, is rapidly consumed by infiltrating immune cells ^83^, suggesting additional potential metabolic constraints inducing antibiotic tolerance ^84^. Testing this idea would require methods to quantify local arginine concentrations around extra- and intracellular *S. aureus*.

*S. aureus* down-regulated the main peroxide-detoxifying enzyme catalase KatA ^85^ and superoxide dismutase SodA ^86^, and maintained unaltered levels of NO-detoxifying flavohemoprotein Hmp ^87^ (Fig. 3a; Ext Data Fig. 2a). This could reflect limited oxidative and nitrosative attacks by human phagocytes at low oxygen ^72^ and aligns with the dispensability of KatA for infections in mice ^76,88^. Unaltered levels of the defense protein DltA (Ext Data Fig. 2a) suggested limited stress by host antimicrobial peptides ^89^.

Overall, *S. aureus* appeared mainly limited by metabolic factors. To test this idea, we prepared fresh, unfixed biopsy slices. Initially, the slices contained mainly isolated *S. aureus* cells as observed before (Extended Data Fig. 2d). After 16h in glucose- and amino acid-rich, oxygenated conditions, large clusters with active cell-wall synthesis appeared, indicating vigorous replication dynamics (Fig. 3c; Extended Data Fig. 2d) and resolved metabolic constraints. However, this approach succeeded for only one biopsy, as other unfixed live biopsies were too soft for slicing.

### Recapitulating in vivo conditions explains poor antimicrobial efficacy

Our analysis revealed that *S. aureus* encountered diverse conditions in musculoskeletal infections, some of which may contribute to the poor clinical efficacy of antibiotics. To test this idea, we evaluated several conditions individually in simplified in vitro assays using genetically diverse clinical isolates.

To test metabolic constraints affecting extracellular bacteria, we used a human plasma-like medium (Plasmax) ^90^ without glucose and 11 mM lactate at 0.5% O₂ and mildly acidic pH 6.6 (often associated with inflammation ^91^). Under these conditions, *S. aureus* replicated slowly (Fig. 4a). The two first-line β-lactam antibiotics flucloxacillin and cefazoline (but not the fluoroquinolone levofloxacin) showed reduced activity (Fig. 4b), consistent with growth-rate-dependent β-lactam activity ^92^. Thus, metabolic constraints present in infected tissues may impair antibiotic efficacy.

**Figure 4:**
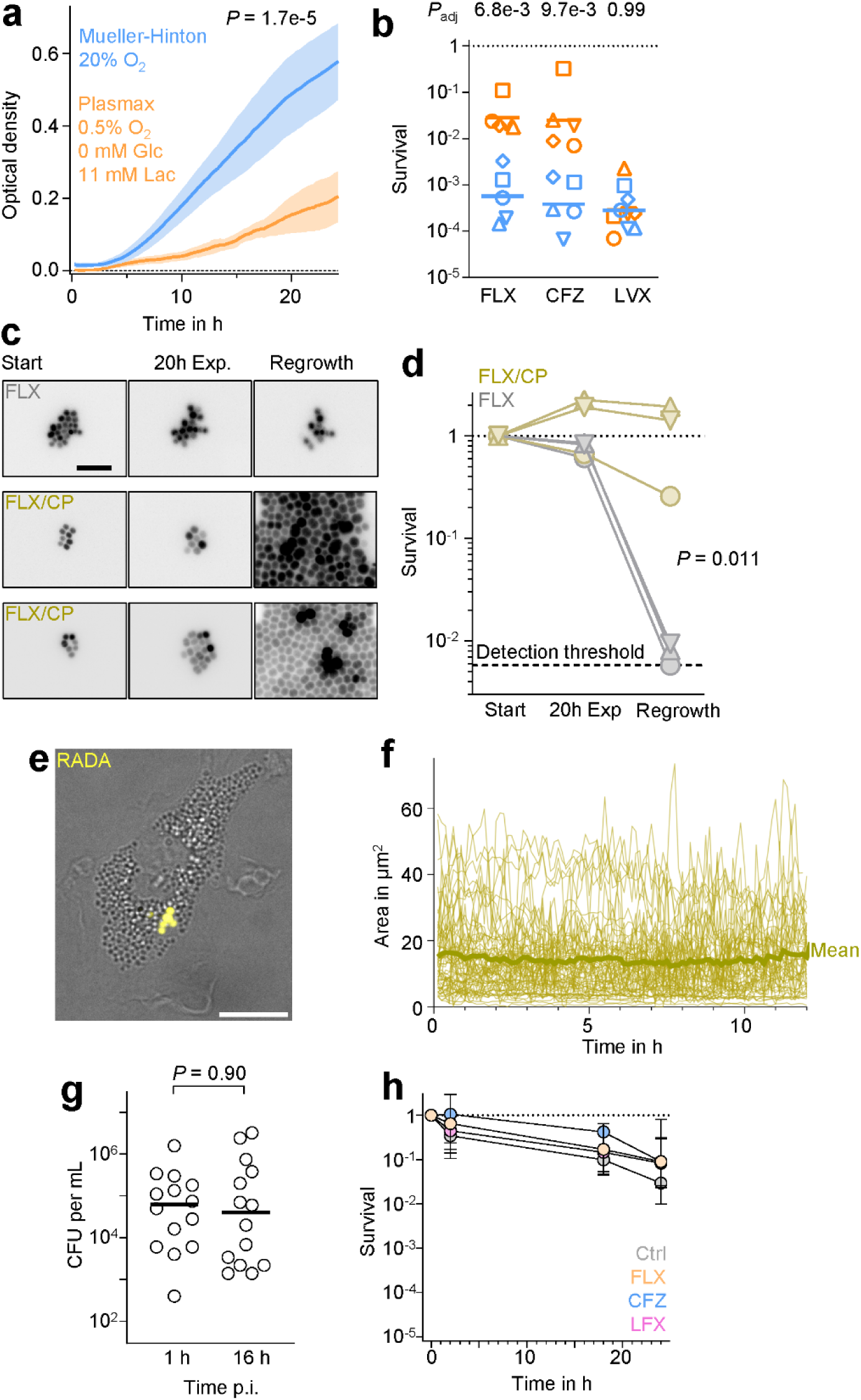
*S. aureus* resilience to antibiotics in patient-mimicking conditions. **(a)** Growth of *S. aureus* clinical isolates in multi-well plates in standard Mueller-Hinton broth or tissue-mimicking conditions (glc, glucose; lac, lactate). Shown are means and SDs for 5 *S. aureus* strains (each an average of 4 technical replicates). One-way ANOVA testing the difference between conditions. **(b)** Survival of *S. aureus* clinical isolates in axenic microaerophilic growth conditions after 24 h exposure to flucloxacillin (FLX), cefazolin (CFZ), or levofloxacin (LVX) in Mueller-Hinton broth or tissue-mimicking conditions. All strains had the same MICs for FLX (0.3 mg/L) and LVX (0.2 mg/L). Each symbol represents a different clinical isolate. Each data point represents the geometric mean of 4 to 6 independent experiments. Horizontal bars represent geometric means. Paired t-test of log-transformed data with correction for multiple comparisons according to Holm-Šídák (*P*_adj_, adjusted *P*-values). **(c)** Responses of GFP-*S. aureus* to flucloxacillin (FLX) with or without calprotectin (CP). Shown are inverted fluorescence signals in greyscale before (Start) and after 20h flucloxacillin exposure (20h Exp.) as well as after 6h regrowth after flucloxacillin wash-out. Scale bar, 5 μm. **(d)** Quantification of data shown in panel c (Start, start of flucloxacillin treatment; 20h Exp, 20h exposure). Shown are results for three different strains with each 93 to 322 individual cells per strain. Paired t-test of log-transformed regrowing fractions. **(e)** Infected monocyte-derived macrophage containing *S. aureus* labelled prior to infection with RADA. Scale bar, 10 μm. **(f)** Area of intracellular *S. aureus* during 12 h in monocyte-derived macrophages. Each line represents one macrophage. The thick line represent the mean. **(g)** Survival of intracellular *S. aureus* at 1 h and 16 h post-infection (p.i.). Each circle represents a different well. The horizontal bars represent geometric means. t-test of log-transformed data. **(h)** Survival of intracellular *S. aureus* during exposure to flucloxacillin (FLX), cefazolin (CFZ), or levofloxacin (LVX), or under control conditions (Ctrl). Geometric means and geometric SDs of three clinical isolates are shown.

Extracellular *S. aureus* were often entangled in extracellular traps containing calprotectin (Fig. 2f, g), which is elevated during musculoskeletal infections ^93^. Calprotectin sequesters transition metals ^94,95^ and may affect antibiotic activity, though prior results are contradictory ^96–98^. In microfluidic devices with a low-nutrient medium that supported ∼1h generation time, flucloxacillin stopped *S. aureus* growth but induced little lysis (as detected by loss of cytosolic GFP) (Fig. 4c, d), consistent with reduced β-lactam activity at low metabolic rates ^99^. After drug washout, almost no cells resumed growth within 6h, indicating an extended post-antibiotic effect. Addition of 120 mg/mL ^96^ calprotectin had no significant impact on pre-treatment growth but permitted some growth during exposure and largely abolished the post-exposure replication arrest, resulting in vigorous proliferation (Fig. 4c, d). A recent preprint reported also calprotectin inhibition of β-lactam activity ^100^. Thus, calprotectin surrounding *S. aureus* in infected tissues may reduce antibiotic efficacy.

To assess the impact of intracellular location, we infected primary monocyte-derived macrophages (CD14⁺ CD16⁺ CD68⁺) (Fig. 4e), approximating the main infected host cell type in biopsies. After phagocytosis, most *S. aureus* survived for at least 16h without replicating (Fig. 4f) ^101^, consistent with the predominance of intracellular singlets in biopsies (Fig. 2i, j). Some infected cells lysed, releasing bacteria to the extracellular space. Several clinically relevant antibiotics showed poor activity against intracellular *S. aureus* (Fig. 4g), consistent with prior work in other cell types ^102–107^. Thus, the dominant intracellular residence in CD14⁺ CD16⁺ CD68⁺ cells in infected tissues may contribute to poor antibiotic efficacy.

Thus, various conditions experienced by *S. aureus* in human tissues independently reduced antibiotic efficacy. Assays employing these conditions might enable physiologically relevant antimicrobial susceptibility testing ^108^ for developing new control strategies.

## Discussion

The replication rates, aggregation state, and host-cell context of bacterial pathogens within infected tissues are critical determinants of disease progression, therapy, and prevention, but data from human infections are scarse. Here, we addressed this gap for *S. aureus* musculoskeletal infections.

To cover a broad spectrum of clinical presentations, we analyzed a heterogeneous patient cohort, multiple tissue types, and genetically diverse *S. aureus* strains. Despite this diversity, common features of *S. aureus* infection biology in human tissues emerged. The majority of *S. aureus* resided intracellularly, challenging the model of predominantly extracellular pathogenesis with limited intracellular states ^5,109^. The dominant infected host cell type, non-classical monocytes/macrophages, is abundant in chronically inflamed tissues ^40,41,110,111^ and in *S. aureus*-infected non-human primates ^112^, yet remains underexplored in bacterial pathogenesis. These findings highlight a physiologically relevant but insufficiently studied niche for future mechanistic work. Moreover, most *S. aureus* were single cells or pairs with apparently limited replication and high antibiotic tolerance, challenging models that attribute antibiotic resilience primarily to large clusters and biofilms ^5,49^. Nonetheless, large clusters/biofilms might be prominent on bone and implant surfaces, which we did not examine.

Based on our data, we propose different stages of disease. During early infection, *S. aureus* likely undergoes rapid aerobic growth ^113^, with increasing bacterial loads triggering escalating inflammation. In advanced disease stages requiring surgery, host inflammation appeared to suppress *S. aureus* replication. Infiltrating non-classical monocytes/macrophages phagocytosed *S. aureus* and restricted intracellular growth. However, *S. aureus* retained the capacity to express virulence factors, including toxins that lysed infected host cells, releasing bacteria into the extracellular space. *S. aureus* replication was restricted by low oxygen and reduced glucose availability due to impaired tissue perfusion and inflammation-associated metabolic shifts. The overall slow *S. aureus* growth likely contributed to poor antibiotic efficacy ^114–116^. Antibiotics active against non-/slowly growing bacteria (e.g., ADEPS ^117^) and combinations of anti-virulence strategies with antibiotics represent promising treatment avenues ^118–120^. Physiologically relevant in vitro models that recapitulate in-patient conditions ^4,121,122^ will be key to advancing these strategies.

Several vaccines candidates relying on antibody-mediated immunity failed in clinical trials ^18,122,123^. *S. aureus* vaccine antigens (protein A, α-hemolysin, IsdB, MntC) were detectable in biopsies (Extended Data Fig. 2a), arguing against insufficient antigen expression as the primary explanation. Instead, the predominance of intracellular *S. aureus* likely limited direct antibody access. Antibodies may still opsonize extracellular for phagocytic killing ^124^, but phagocytes in human tissues showed limited killing of intracellular bacteria *S. aureus* (Fig. 2b) ^125^. Tissue-adapted bacteria might be less vulnerable to phagocyte killing due to enhanced toxin expression Fig. 3a, than in vitro-grown *S. aureus* used in standard opsonophagocytosis assays. Effective protection might require engaging multiple arms of immunity, including cellular immunity, toxin neutralization, and strategies to overcome non-protective immune imprints from prior exposure ^18,123,126–128^.

Our study has limitations. The patient cohort was single-center with single time points. The study was underpowered to relate findings to different tissue types, antibiotic regimens, or outcomes. We did not include bone, implants, or synovial fluid, which may harbor biofilms. Vancomycin staining cannot distinguish different Gram-positive bacteria, restricting analysis to mono-microbial *S. aureus* infections. Infected host cell types were characterized based on few markers. Finally, proteomics measured bulk protein content without distinguishing intra- and extracellular bacteria.

In conclusion, multidisciplinary collaboration, AI-guided imaging, and ultrasensitive proteomics revealed fundamental features of *S. aureus* infection biology directly in human tissues and established physiologically relevant in vitro models. Extending such multi-patient, spatially resolved datasets to other bacterial infections will be crucial for accelerating the development of mechanism-based prevention and treatment strategies.

## Methods

### Patient sampling and ethics

Research performed on samples of human origin was conducted with approval of the local Ethical Review Board (Ethikkommission Nordwestschweiz, Project-ID 2020-02588) and performed in compliance with all Swiss and local laws and ordinances. De-identified human tissue samples were collected after written informed consent was acquired. Tissue samples from *S. aureus* infections were collected intraoperatively during revision surgeries for musculoskeletal infections performed between 2017 and 2024 at the University Hospital of Basel, Switzerland. Sampling was restricted to debrided areas that were macroscopically infected, and only tissues that would be usually discarded were used for research. Samples consisted of infected soft tissue, joint tissue, or infectious tissue and membranes adjacent to affected implants or bones. For validation of imaging data, FFPE tissue blocks from routinely retrieved histology samples from 14 additional individuals were also included (Extended Data Table 1).

### Treatment, sectioning, and staining of live biopsies for imaging (cohort 1)

Intraoperatively collected biopsies were collected in pre-cooled 50 mL tube without buffer on ice and transferred to the lab. Using a puncher, cylindrical pieces with 6 mm were prepared and embedded in 8% low gelling-temperature agarose (Roth, 6351.5) followed by rapid cooling in a 4°C fridge. Tissues were cut into 250 μm sections using a Compresstome and transferred to a 24-well tissue culture plate containing 300 μL RPMI containing 10% human serum, 1% glutamax, and 1% non-essential amino acids. Sections were maintained at 37°C and 5% CO_2_ on a rotatory shaker. Some sections were immediately stained with 1 mM RADA (an orange-red TAMRA-based fluorescent D-amino acid; Tocris, 6649), 5 mg/L Hoechst 33342, and 0.25 mg/L Vancomycin-Janelia Fluor 669 (Van-JF669) ^28^ for 30 min at 37°C, washed thrice with PBS, fixed with 4% paraformaldehyde at room temperature, washed with PBS, and mounted in 40µL mounting medium (Agilent DAKO, Cat No. S3023). Other sections were incubated in tissue culture medium for 16h and then labeled with RADA, Hoechst and Van-JF669 as described above.

### Treatment, sectioning, and staining of fixed biopsies for imaging (cohort 1)

Intraoperatively collected biopsies were immediately fixed in 4% paraformaldehyde for 48h at 4°C. Samples were then put into cryoprotectant buffer (300 g sucrose, 10 g polyvinyl-pyrrolidone 40, 400 mL ethylene glycol, 400 mL PB buffer (10 mM NaH_2_PO_4_, 40 mM Na_2_HPO_4_). Once the tissue had sunk to the bottom, the sample was stored for at least one week up to several years at -20°C to reduce tissue autofluorescence ^27^.

For sectioning, stored tissue was cut into several pieces with a disposable scalpel, washed thrice with phosphate-buffered saline (PBS, Life Technologies, 20012019) for 5 min at room temperature, and embedded in 8% agarose (Sigma, A6013). Tissues were cut into 60 μm sections using a Compresstome (VF-310-0Z, Precisionary Instruments). Pools of five adjacent sections were stored together in cryoprotectant at -20°C.

The 60 μm thick sections would yield non-interpretable images with standard hematoxylin and eosin (H&E) staining due to large overlap of the tissue layers. To circumvent this issue, we adopted a virtual hematoxylin and eosin approach based on nuclear staining and tissue autofluorescence ^33^. Sections were brought to room temperature, washed thrice with PBS, and incubated in PBS containing 0.02% sodium azide (to prevent microbial contamination) at 4°C for 1 week to restore autofluorescence. Sections were washed thrice with Tris-buffered saline pH 7.4 (TBS) at room temperature. After staining with 1mg/L 4’,6-diamidino-2-phenylindole (DAPI) at room temperature for 20 minutes, sections were washed thrice with TBS and mounted in 40µL mounting medium (Agilent DAKO, Cat No. S3023).

To stain bacteria and various host components, sections were washed thrice for 5 min with TBS at room temperature. For antigen retrieval, sections were incubated in 1 mL prewarmed 10 mM sodium citrate, pH 8.5, at 37 °C for 30 min. Sections were permeabilized by washing thrice with TBST (0.1% Triton X-100 in TBS), blocked for 30min with blocking buffer (1% BSA-fraction V, 2% human serum in TBST), and washed thrice with TBST. Endogenous biotin and biotin-binding sites were blocked using the streptavidin/biotin blocking kit (Vector Laboratories, VC-SP-2002) following the manufacturer’s protocol. Sections were incubated in primary antibodies (Extended Data Table 2) in blocking buffer at 4°C overnight, washed thrice with TBST, and stained with secondary antibodies (Extended Data Table 2) for 1h at room temperature. Sections were washed thrice with TBS and incubated for 10 min with 0.25mg/L Vancomycin-Janelia Fluor 669 ^28^ and in some cases 1 mg/L DAPI. Sections were washed thrice with TBS and mounted in 40µL mounting medium (Agilent DAKO, Cat No. S3023).

Human Fcγ RIII (CD16) Antibody was biotinylated using Mix-n-Stain™ Biotin Antibody Labeling Kit (Biotium, 92444) according to the manufacturer’s protocol and detected with Streptavidin-Alexa Fluor 488 (Invitrogen, S11223) or Streptavidin-Horseradish Peroxidase (Invitrogen, S911) with CF 405S Tyramide (Biotium, 92917).

For staining *S. aureus* 16S rRNA, tissue sections were washed with RNase-free TBS and placed in 50% ethanol for 3h at -20°C to permeabilize the tissues ^37^. After washing twice with TBS, *S. aureus* cell wall was digested with 1g/L lysozyme in 10 mM Tris-HCl pH 8.0 for 10 min at 30°C. After washing twice with TBS, sections were incubated in pre-warmed hybridization buffer (2 M Urea, 4 M NaCl, 50 mM Tris–HCl, 1g/L BSA, 250mg/L yeast tRNA, 500 mg/L Salmon sperm DNA pre-heated to 95°C, pH 7.5) containing 400 nM LNA/2’OMe probes (sense probe, 5’ TAMRA - [A]{G}[C-A]{A}[G-C]{T}[U-C]{T}[C-G]{T}[C-C]; or control probe, 5’ TAMRA - [U]{C}[G-U]{T}[C-G]{A}[A-G]{A}[G-C]{A}[G-G]) and incubated for 1.5h at 65°C in humidified chamber. Tissues were washed with 5 mM Tris-base (pH 10), 15 mM NaCl, and 0.1% (v/v) Triton X-100 and incubated in the same solution for 30 min at 65°C. Afterwards, immunostaining and labeling of DNA and *S. aureus* was done as described above, except using a higher Van-JF669 concentration (0.5 mg/L).

### Sectioning and staining of formalin-fixed paraffin sections (cohort 2)

Formalin-fixed paraffin-embedded (FFPE) musculoskeletal tissue samples were cut into 4 μm sections, deparaffinized with two 10 min washes in 100% xylene at room temperature, followed by two 5 min incubations in 100% ethanol, 2 min each in 96%, 70%, and 50% ethanol, and distilled water. For antigen retrieval, tissues were incubated at 95°C for 20 min in EDTA-based epitope retrieval buffer pH 9.0 (Biosystems, AR9640,), then cooled for 15 min at room temperature. Slides were transferred to PBST (PBS + 0.1% Tween 20) for permeabilization. Tissue structure was circled with a hydrophobic pen (Abcam, ab2601), 250 µL blocking buffer (PBST with 5% goat serum) was added, and incubated for 1h in a dark humidity chamber. Blocking buffer was removed by tilting, and sections were stained with primary antibodies (Extended Data Table 2) in blocking buffer overnight at 4°C in the dark. Primary antibodies were removed by tilting and rinsing thrice with PBST. Sections were stained with secondary antibodies (Extended Data Table 2) for 1h at room temperature in the dark. Sections were washed thrice with PBST and thrice with PBS. Sections were stained with 1.7 mg/L Van-JF669 ^28^ and 1.25 mg/L DAPI (Abcam, ab228549) in PBS for 15 min at room temperature in the dark. Slides were rinsed thrice in PBS and mounted with two drops of aqueous mounting medium (Merck, F4680) and covered with 0.15 mm glass coverslips.

### Microscopy and image analysis for vibratome sections (cohort 1)

Tissue sections were imaged using an Evident/Olympus spinning disk IXplore SpinSR system with Yokogawa CSU-W1 confocal scan head (50 µm disk or Sora disk) equipped with 405 nm, 488 nm, 561 nm, and 640 nm diode lasers, emission filters (422-472 nm, 500-550 nm, 580-653 nm, 665-705 nm), a 561 nm long-pass beam splitter for dual-camera mode, and Hamamatsu ORCA-Fusion sCMOS cameras.

Image stacks of sections stained for virtual hematoxylin and eosin were acquired using an UPL X APO 10x 0.4 NA air objective, a z step size of 2 μm, 405 nm excitation (30% laser power)/422-472 nm emission and 488 nm excitation (40% power)/500-550 nm emission, and 200 ms exposure times. Images were analyzed using Fiji v1.54f ^129^. The middle plane of the image stack was median filtered (radius 2). Fluorescence intensities in both channels were log-transformed ^33^ and contrast-adjusted to cover the upper half of the intensity scale (lowest pixel value at midpoint of scale; highest pixel value at maximum of scale). The DAPI channel was set to yellow, the green autofluorescence channel to cyan. The two-channel image was then transformed to RGB and inverted to yield images resembling standard histology H&E staining.

For feedback (smart) imaging of sections stained for *S. aureus*, DNA, 16S rRNA, and host components, ScanR acquisition software (version 3.3.0 Evident/Olympus) was used. Overview pre-scan tiled acquisitions were done using a UPL APO 60x 1.5 NA oil objective with a voxel size of 108 x 108 x 300 nm³ covering the top 25 µm of the 60 µm section (84 z planes) and dual-camera mode using channels 488 nm excitation/500-550 nm emission and 640 nm excitation/665-705 nm emission (laser powers were adjusted once for each section to reach the linear range of the cameras), and 200 ms exposure times.

Z stacks were automatically converted to maximum intensity projections. These projections were analyzed during the acquisition of the next tiles with a specifically trained semantic Neural Network (NN) to segment Van-JF669 positive bacteria. To train the NN, we created a training dataset based on real images analyzed with classical methods using manual thresholds (elongation, Max_intensity 640nm_zMax, mean intensity ratio between main object and periphery, area of object). We manually curated the detected objects in three classes (positive, Extended Data Fig. 1d; negative; and ignore for unclear cases). This process was replicated on two samples yielding 484 positive objects (single bacteria or clusters). The *positive* detection class was used to train a dedicated NN infrastructure provided by the acquisition software with 50,000 iterations. Validation of the NN segmentation on 18 additional sections from 9 patients, yielded 56,827 segmented objects with 26,773 true positive, 20,035 false positive, and 1,255 false negative (4.7% of all bacterial objects). The false positives were mostly tissue fibers (Extended Data Fig. 1c, cyan arrowheads).

Each tile with NN-detected bacteria in the center region matching the field of view of the 100x objective was immediately rescanned using an UPL APO 100x 1.5 NA oil objective with a voxel size of 65 x 65 x 210 nm³ covering the top 25 µm of the 60 µm section (119 z planes). We used dual-camera mode with channels 405 nm excitation/422-472 nm emission; 561 nm excitation/580-653 nm emission and 488 nm excitation/500-550 nm emission; 640 nm excitation/665-705 nm emission (laser powers were adjusted once for each section to reach the linear range of the cameras; except for rRNA quantification where laser power was kept at 30%), and 200 ms exposure times. The complete acquisition run duration (60x scan, image analysis and 100x rescan) required between 6 and 12 hours per mm^3^ tissue.

NN-segmented *S. aureus* were manually validated in the pre-scan 60x acquisition max-projections and used for tissue-wide bacterial density maps. NN-segmented and validated *S. aureus* in the high-resolution 100x z-stacks were manually quantified (number of bacteria in a cluster) and classified intra- and extracellular (based on a channel combining antibodies against seven human cytosolic antigens: ALDOA, ENO1, GAPDH, HADH, PDAP1, RPL10, RPL17; Extended Data Table 2) as well as infected human cell type using Fiji v1.54f ^129^. When more than 30 100x Z-stacks were acquired, 30% of the acquired data were randomly selected for analysis.

For super-resolution imaging 100x 3D z-stacks (step size = 130 nm) were acquired using the super resolution module in CellSens software version 4.1.1 (SoRa disk and 3.2x magnification). 405 nm and 640 nm lasers were used at 30% and 50% power with 200 ms exposure times. Raw images were deconvolved using Huygens professional 21.10.1p2 64b (Scientific Volume Imaging, The Netherlands, http://svi.nl) with following parameters: Classical Maximum Likelihood Estimation (CMLE) algorithm, 25 iterations, 0.01 quality change, signal ratio = 30.6 (405=DAPI) and 23.6 (640=Van-JF669), back projected pinhole radius = 78 nm and back projected pinhole spacing = 2.53 µm.

### Microscopy and image analysis for paraffin sections (cohort 2, validation)

Tissue sections were imaged using a Nikon Ti2-E inverted microscope with a spinning disk unit (Crest V3, CrestOptics, pinhole size = 50 µm) and Plan Apo Lambda S 40x NA 1.25 silicone and Plan Apo Lambda S 100x NA 1.35 silicone objectives. Fluorescence was excited with a Celesta light engine with solid-state lasers (Lumencor) at 405, 477, 546, 640, 749 nm and collected with a penta-edge 421/491/567/659/776 nm dichroic beam splitter, a dual-edge 471/539 nm splitter, and 438/24 nm (DAPI), 511/20 nm (tissue autofluorescence), 560/25 nm (Alexa Fluor 555), 685/40 nm (JF669) single-bandpass filters and a 441/30, 511/26, 593/36, 684/34, 817/66 nm (Alexa Fluor 750) penta-bandpass filter. Images were acquired with a Photometrics Kinetix camera controlled with the Nikon NIS software. Excitation light intensity and camera exposure were adjusted for each channel to prevent image oversaturation and were kept the same for samples from the same patients.

The NIS JOBS module was used for automated image acquisition. First, a tiled overview image (15% tile-overlap) of the entire tissue section was captured using the 40x objective for all channels for a total of 9 z-planes with 0.5 μm z-step size. The Nikon Perfect Focus System was used to keep the focus over large areas. Subsequently, maximum intensity projections of the tiles were stitched for identification of bacterial objects. The NIS GA3 module was used for object identification, which was performed as follows: the Gaussian blur (sigma = 1) of the autofluorescence channel was subtracted from the JF669 channel to obtain a background corrected image of the bacteria. Then a threshold manually adjusted per image in the range from 3 - 10 for the JF669 channel defined objects of interest. Tiles with objects of interest in its center were rescanned using the 100x objective and 13 z-planes with 0.3 µm step size around the focus plane detected by auto-focusing on the JF669 channel.

Analysis of 100X high-resolution 3D images were performed in ImageJ, 2.14.0 ^130^, NIS-Elements AR Analysis 5.42.06 (Nikon), or directly on the data storage platform OMERO ^131^ (hosted by the IMCF, Biozentrum University of Basel). In tissues with high bacteria abundance, at least 100 bacteria per tissue were classified. In tissues with low abundance, all bacteria were classified. Bacteria were segmented by a pixel classifier which was trained on randomly selected images on the JF669 channel and manually validated. For validated objects, we counted the number of bacteria, determined their DNA content, and classified them as intracellular or extracellular based on enclosure by vimentin-stained human cell boundaries. For intracellular bacteria, the CD45 status of the human host cell was also assessed.

### Treatment and staining of fresh unfixed biopsies for flow cytometry

Biopsies were placed in RPMI 1640 + 10% human serum + 2mM EDTA and single-cell suspensions were obtained using the Multi Tissue Dissociation Kit (Miltenyi Biotec) in combination with the gentleMACS Dissociator (Miltenyi Biotec) according to the manufacturer’s protocols with some modifications. Biopsies were minced and mixed with the enzymes “D”, “R” and “A” in RPMI 1640 and digested using the gentleMACS program mr_SMDK_1 at 37°C for 60 min. Single-cell suspension was filtered using 40 μm strainers and mononuclear cells were isolated using a Percoll-based density gradient centrifugation. Red blood cells (RBC) were lysed using a RBC lysis buffer (eBioscience). Remaining cells were incubated with FcBlock (TruStain FcX) and stained with appropriate surface markers. Cells were fixed with 4% paraformaldehyde. Cells were analyzed using Cytek® Aurora spectral flow cytometer and FlowJo (BD, version 10.8.1). The gating strategy is show in Supplementary Figure 1.

### Whole-genome sequencing of *S. aureus* isolates

Freshly frozen tissue biopsies were thawed and plated on blood agar. Colonies were verified as *S. aureus* by sequencing the 16S rRNA gene after PCR-amplification using primers AGAGTTTGATCCTGGCTCAG and CGGTTACCTTGTTACGACTT. Oxford Nanopore long-reads and Illumina short reads were obtained by SeqCenter (Pittsburgh, USA). Short reads were trimmed using TRIMMOMATIC/trimmomatic-0.39.jar ^132^. Hybrid assembly was done with Unicycler/v0.5.1 ^133^ and assembled using Flye (2.9.4-GCC-13.2.0) and annotated using Prokka (1.12-goolf-1.7.20-BioPerl-1.7.2, including ncDNA and rRNA predictions using Barrnap, HMMER and Infernal) ^134^. Clonal complexes and sequence types were identified using the PubMLST database (https://pubmlst.org/) ^135^.

### Auxotrophy and nutrient utilization of *S. aureus* strains

*S. aureus* was grown in chemically defined Hussain-Hastings-White medium (HHW) ^136^. For identifying amino acid auxotrophies (Extended Data Fig. 2c), one amino acid was left out and 4.6 mM D-gluconate was added as carbon/energy source. Growth was monitored for 24h in a microplate spectrophotometer (BioTek, Epoch 2) and OD_600 nm_ values at 24h were normalized by values obtained in full HHW with all amino acids.

For quantifying nutrient utilization (Extended Data Fig. 2b), we used HHW with reduced concentrations of those amino acids that *S. aureus* can utilized as carbon sources ^137^, to minimize baseline growth without added carbon/energy source (alanine, 0.11 mM; arginine, 0.06 mM; aspartate, 0.11 mM; glutamate, 0.1 mM; glycine, 0.2 mM; histidine, 0.06 mM; threonine, 0.13 mM; proline, 0.13 mM; serine, 0.1 mM; tryptophan, 0.05 mM). Various nutrients were added at 0.1% (w/v) and growth was monitored for 24h in a microplate spectrophotometer (BioTek, Epoch 2). OD_600 nm_ values at 24h were normalized by values obtained for each strain with the respective best nutrient and in absence of added nutrient.

### Mass spectrometry-based proteomics of unfixed biopsies

Intraoperatively collected biopsies were immediately frozen at -20°C in the operation room and later transferred to a -80°C freezer for storage. The tissues were thawed on ice, 10 mL ice-cold PBS was added, and the tissue was homogenized in gentleMACS M tubes in a gentleMACS dissociator (Miltenyi) using the mouse spleen program. After centrifugation at 16,000xg and 4°C for 5 min, the pellet was resuspended in 2 mL hypo-osmolar buffer (0.1% Triton-X, 5 mM EDTA) to deplete host cells and disrupted again in the gentleMACS dissociator. After centrifugation at 8,000xg and 4°C for 10 min, the pellet was resuspended in 2 mL mammalian lysis buffer (50 mM Tris-HCl, 5 mM EDTA, 0.5% NP-40, 100 mM NaCl, pH 8.5) followed by another disruption in the gentleMACS dissociator, and centrifugation at 8,000xg and 4°C for 10 min. The pellets were stored at -80°C until further analysis. Recovery of *S. aureus* was followed by quantitative plating. The overall bacterial yield was usually 20-50%. *S. aureus* was confirmed by PCR-amplification and sequencing of the 16S rRNA gene as described above.

*S. aureus* proteins were detected in the final pellets by targeted mass spectrometry in biopsy homogenates using the SureQuant approach as recently described ^58^. Data were analyzed as described and normalized based on signal intensities for proteins DnaK (Uniprot identifier Q2FXZ2) and DNA-binding protein HU (Uniprot identifier Q2FYG2). Statistical analysis was performed using Perseus (1.6.14.0) ^138^. The calculated false discovery rate q was based on data permutations and significance-analysis-of-microarrays statistics using a s0 value of 0.1. Depending on sample availability, we analyzed 7 or 8 out of a total of 12 different biopsies. We show data for proteins detected in at least three of the analyzed biopsies.

### Quantification of glucose and lactate in biopsies

Intraoperatively collected biopsies were immediately frozen at -20°C in the operation room and later transferred to a -80°C freezer for storage. The tissues were thawed on ice, homogenized, and de-proteinized using 10 kDa spin columns (Abcam, ab93349), and glucose and lactate were quantified using enzymatic assays (Glucose detection kit, Abcam, ab102517; L-lactate assay kit, Abcam, ab65331) according to the manufacture’s protocols.

### Growth and antibiotic exposure under hypoxia

*S. aureus* was grown in glucose-free Plasmax cell culture medium (CancerTools.org, 161587) containing 11 mM lactate and adjusted to pH 6.6 in a hypoxystation (Whitley, H35) at 0.5% O_2_ / 5% CO_2_ and 37°C. Growth was monitored in a microplate spectrophotometer (BioTek, Epoch 2). For antibiotic exposure, stationary phase cultures grown under the same conditions were diluted 1:100 in pre-warmed fresh medium. Antibiotics (flucloxacillin, 10 mg/L; cefazolin 14 mg/L; levofloxacin, 9 mg/L) were added after 5h of growth in tubes stirred with magnetic bars). Cultures were plated on terrific broth agar before and after 24h of exposure and incubated in the hypoxystation. Strains tested were P25, P28, P62, P70, P128 (Extended Data Table 3).

### Microfluidics experiments

*S. aureus* carrying a plasmid driving constitutive *gfp* expression ^139^were grown overnight in BHI medium, diluted to OD_600_ 0.05 in BHImod medium (1:500 BHI diluted in saline with 1mM MgCl_2_, 2mM CaCl_2_ and 6.2 mM ß-mercaptoethanol), grown for 4h, and inoculated in the microfluidic plates at 37 °C with humidity controlled by Okolab T-unit (Okolab). After growth for 3h in BHImod (generation time ∼1 h) with/without 120 mg/L calprotectin, bacteria were exposed for 20h to 12 mg/L flucloxacillin in the same medium, and then switched to normal BHI without calprotectin.

Bacteria were imaged using a Nikon Eclipse Ti2 inverted microscope equipped with Perfect Focus System and Plan Apo 100× oil Ph3 DM NA 1.45 objective (MRD31905), SPECTRA X light engine (Lumencor), phase contrast (8% light intensity, 200 ms exposure) and a Dualband EGFP/mCherry filter set (Chroma 59022 ET; 470 nm excitation, 4% light intensity, 150 ms exposure), a Hamamatsu ORCA-Flash4.0 V3 CMOS camera (C13440-20CU) with pixel size of 65 nm, and NIS-Elements (Nikon). Images were acquired at 5 min intervals. Images were analysed with Fiji 1.54q ^129^ using plugins MultiStackReg ^140^ and FeatureJ-Laplacian (Smoothing scale 2.0) (developed by E. Meijering). Strains tested were P28, NCCR74, NCCR125 (Extended Data Table 3).

### Cell culture infection with monocyte-derived macrophages

Human monocytes were obtained from anonymous healthy human donors (Blood Donation Center, University Hospital Basel). PBMCs were isolated using ficoll density gradient centrifugation. Monocytes were isolated by negative bead selection using the human pan monocyte isolation kit (Miltenyi Biotec, 130-096-537) according to the manufacturer’s protocol. Monocytes were either directly analyzed by Flow Cytometry or differentiated in 6-well plates at 1x10^6^ per well with RPMI 1640 containing 10% human serum, 10% Glutamax, 10% non-essential amino acids, and 2.5 μg/L M-CSF. Cells were incubated at 37°C with 5% CO_2_. Medium with M-CSF was renewed on day 2. Cells were harvested and plated for infection on day 3.

At the day of infection, plates were washed with PBS and cells were detached by incubation with ice-cold PBS containing 2.5 mM EDTA on ice for 30 min. Detached cells were collected, washed, resuspended in RPMI 1640 containing 10% human serum, and 5,000 cells were seeded black clear-bottom 384-well plates (Perkin Elmer, 6057302) and incubated for 1h at 37°C in a 5% CO_2_ humidified atmosphere. *S. aureus* was cultured overnight in cell culture medium (RPMI 1640 + 10% human serum) at 37°C, 150 rpm. Bacteria were centrifuged at 5,000 rpm for 5 min, washed once with PBS, and stained with 0.25 mM RADA for 15 min at 37°C. Cells were infected with bacteria at a multiplicity of infection of 2-8 bacteria per human cell. After incubation for 1h at 5% CO_2_ and 37°C, extracellular bacteria were removed and the medium was replaced with fresh medium containing 50 mg/L gentamicin or one of the test antibiotics flucloxacillin (32 mg/L), cefazolin (32 mg/L), or levofloxacin (8 mg/L). After various time intervals, the infected cells were washed with PBS and lysed with PBST (PBS containing 0.1% Triton X-100). Serial dilutions of the lysates were plated on Mueller-Hinton agar plates. After overnight incubation at 37°C, colonies were counted. Strains tested were P25, P28, P70 (Extended Data Table 3).

Live-cell microscopy of gentamicin-treated infected cells was performed with a Nikon Ti2-E inverted microscope equipped with a Crest V3 spinning disk unit (CrestOptics) in widefield mode, using either an Apo Plan l 40x NA 0.95 air objective plus the microscope’s 1.5x zoom lens or an Apo Plan l 100x NA 1.45 oil objective (without zoom lens), and the Perfect Focus System for maintenance of the focus over time. Infected cells were kept at 37°C and a 5% CO_2_ humid atmosphere in a H301-K-FRAME stage top incubator (Okolab). RADA was excited with a Celesta Light Engine (Lumencor) at 546 nm and emission was collected through a penta-edge (421/491/567/659/776 nm) dichroic beam splitter and a 595/31 nm single-bandpass filter. Transmitted light was collected with no filter. Images were acquired with a Kinetix camera (Teledyne Photometrics) controlled with the Nikon NIS acquisition software. For each channel, camera exposure was adjusted to prevent over- and underexposure while keeping the laser intensity at 10%. In each well, we acquired 4 non-overlapping fields of view every 7.5 minutes for a duration of 13 hours. Images were analyzed using ImageJ v2.3.0^130^. The original image was duplicated. A max entropy threshold was applied to the RADA channel. Particles of size 0.5μm^3^ were analyzed from the binary image. The masks were applied to the raw image to measure size and mean intensity.

### Statistics

Statistical tests were performed with GraphPad Prism 9.3.1 as indicated in the figure legends. All tests were two-tailed. Proteomics data analysis was performed with Perseus (1.6.14.0). ^138^. The calculated false discovery rate q was based on data permutations and significance-analysis-of-microarrays statistics using a s0 value of 0.1.

## Acknowledgements

We thank Imaging Core and Proteomics Core Facilities, Biozentrum, University of Basel, and the Microscopy Core and Flow Cytometry Core facilities, Department of Biomedicine, University of Basel, for support and technical assistance. We thank Fanny Linder-Hengy, Mandy Mathys, Lea Martin. Kathrin Ullrich, and Nadine Doyle for providing samples to the biobank.

## Author contributions

V.T., B.M., A.A.L., M.A.V., S.J., F.M., N.H., K.S., A.I., A-K.W., J.K., F.E., B.C., S.S., A.E., K.K. performed experiments; V.T., B.M., A.A.L., M.A.V., K.K., T.B., and D.B. analyzed the data; V.T., S.H. and A.F. developed the feedback microscopy workflow for vibratome sections; B.M., and L.S. developed feedback microscopy workflow for FFPE sections; K.J. and P.R.F. developed Van-JF669; R.K., S.M., M.M., and M.C. developed the clinical sampling workflow and collected the intraoperative patient samples; D.Ba. provided expertise on H&E staining; P.M.K. provided clinical microbiology. V.T. and D.B. wrote the paper with inputs from all authors.

## Additional information

Correspondence and requests for materials should be addressed to.

## Data availability

Data points generated for this study are included in the figures whenever possible. All mass spectrometry proteomics data associated with this manuscript have been deposited to the ProteomicsXchange consortium via MassIVE (https://massive.ucsd.edu) with the accession numbers MassIVE MSV000098898 / PRIDE PXD067531. (Dataset is currently private. Reviewers can access the dataset following the instructions on the website. Username: MSV000098898_reviewer, Password: PCF).

**Code availability**: no custom code was used.

---

**Extended Data Figure 1:**
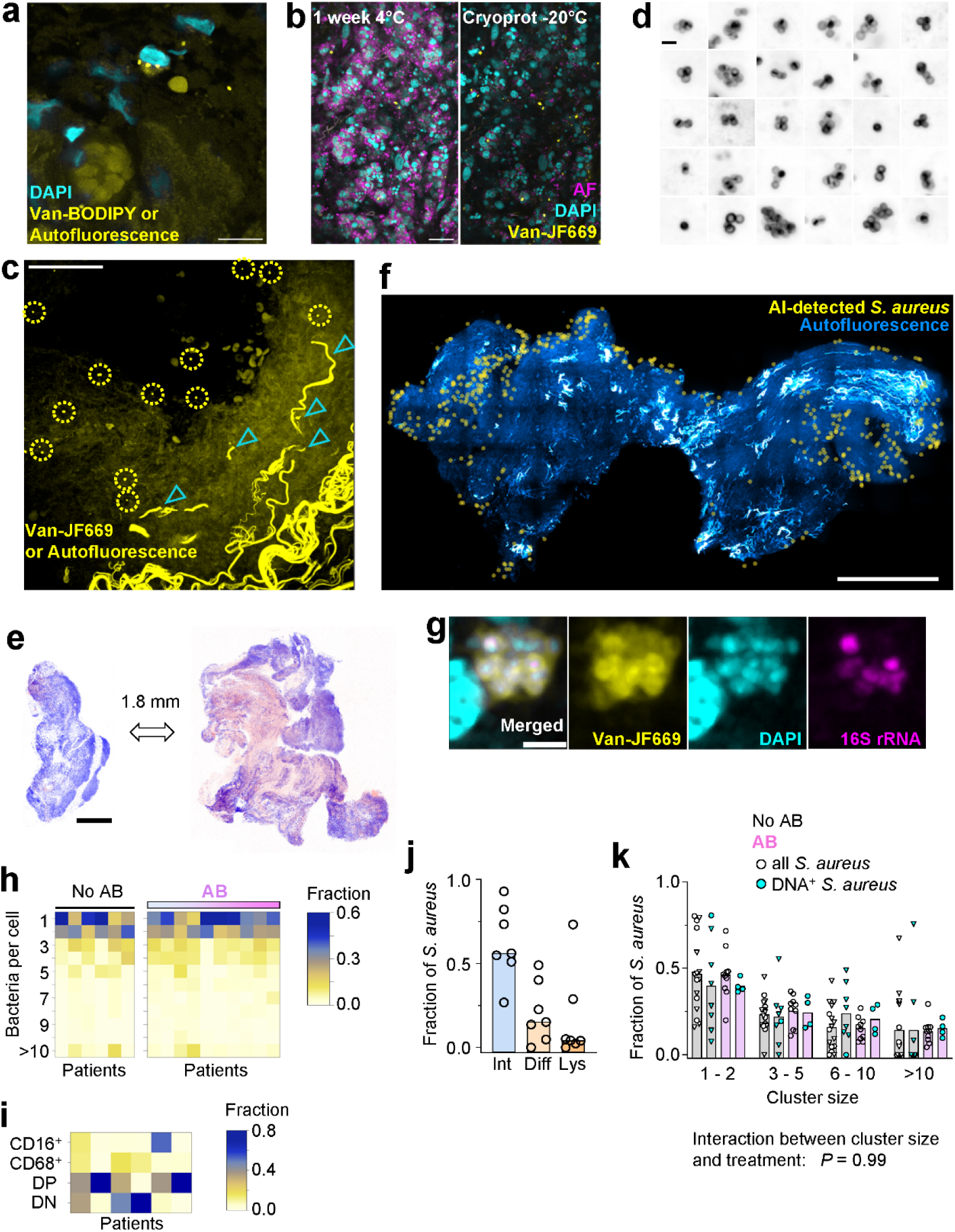
Visualization of *S. aureus* in biopsies. **(a)** Staining of *S. aureus* with commercial vancomycin-BODIPY (Van-BODIPY) and DAPI in a section from a formalin-fixed paraffin-embedded tissue. Diffuse BODIPY-like signals are tissue autofluorescence. Scale bar, 10 μm. **(b)** Impact of low-temperature incubation in cryoprotectant (Cryoprot) on green-yellow fluorescence (shown in false magenta color) in biopsy sections (AF, autofluorescence). Scale bar, 20 μm. **(c)** Fluorescence signals in a vibratome section stained with vancomycin-JF669. The dotted circles indicate objects classified as potential *S. aureus* by a neural network. The cyan arrowheads indicate elastic fibers with broad-spectrum autofluorescence. Scale bar, 50 μm. **(d)** Examples of *S. aureus* appearance in human biopsies. Shown are inverted fluorescence micrographs for improved visibility. Scale bar, 2 μm. **(e)** Two sections from the same biopsy spaced 1.8 mm apart in z direction. The sections were stained with a histology-emulating fluorescence method. Scale bar, 0.5 mm. **(f)** Positions of possible *S. aureus* objects in an overview scan. Scale bar, 0.5 mm. **(g)** Visualization of DNA stained with DAPI and rRNA stained with a fluorescent probe to 16S rRNA in *S. aureus* cells identified with fluorescent vancomycin (Van-JF669) in a formalin-fixed paraffin-embedded tissue section. Scale bar, 2 μm. **(h)** Distribution of intracellular *S. aureus* loads in host cells. Each column represents one patient. Patients receiving antibiotics (AB) are sorted according to length of prior treatment. **(i)** Cell type distribution among CD45^+^ immune cells in biopsies. Each column represents one patient (DP, double-positive; DN, double-negative). **(j)** Distribution of *S. aureus* in host cells with intact nuclei (Int) and diffuse nuclear borders (Diff), or lysed host cells (Lys). Each circle represents one patient; the bars show medians. **(k)** Distribution of DNA-containing *S. aureus* across various cluster sizes in patients treated with antibiotics (AB) or not. Each circle represents one patient (circles, cohort 1; triangles, cohort 2); the bars show means. Two-way ANOVA.

**Extended Data Figure 2:**
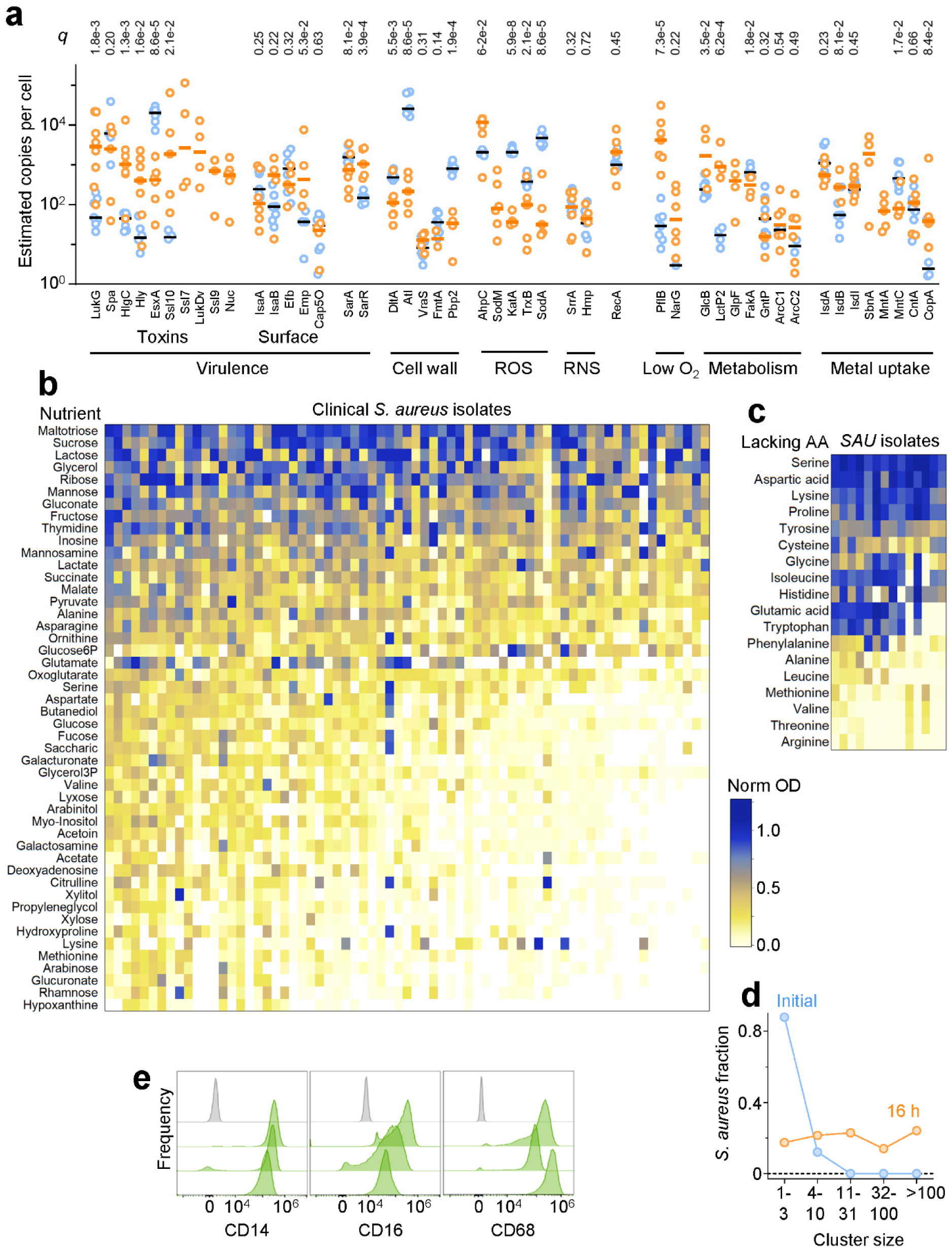
Visualization of *S. aureus* in biopsies. **(a)** Protein abundance in biopsies (orange) and in vitro cultures (blue). Each circles represents a different biopsy or in vitro culture. The horizontal bars show medians. t-test of log-transformed data with calculation of false discovery rates (*q*) according to Benjamini, Krieger, and Yekutieli. **(b)** Growth of clinical *S. aureus* isolates on minimal media containing one main carbon/energy source (Norm OD, normalized optical density after 24 h based on maximal growth with the best respective nutrient and no added nutrient). Each column represents one strain. **(c)** Growth of clinical *S. aureus* isolates on minimal media lacking one amino acid (Norm OD, normalized optical density after 24 h based on maximal growth with the best respective nutrient and no added nutrient). Each column represents one strain. **(d)** Distribution of *S. aureus* in live biopsy slices before (Initial) and after 16 h slice incubation in rich medium at normoxia. **(e)** Expression of three different cell markers in monocyte-derived macrophages. The grey histograms represent isotype control staining. Each green histogram represents a cell population from a different donor.

