## Extended Data Table 1 for "Microgeography of staphyloccoci in human tissue explains antibiotic failure"

| Characteristics | Microscopy cohort 1 (n = 19) | Microscopy cohort 2 (n = 14) | Proteomics (n=12) | Flow cytometry (n=7) | Lactate measurement (n=10) |
|---|---|---|---|---|---|
| Age, years median (IQR) | 65 (52-70) | 64 (56-70) | 68 (61-73) | 69 (56-75) | 73 (58-82) |
| Female, n (%) | 8 (42) | 5 (36) | 1 (8) | 2 (29) | 3 (30) |
| Diagnosis, n (%) | | | | | |
| Prosthetic joint infection | 6 (32) | 8 (57) | 2 (17) | 3 (43) | 7 (70) |
| Fracture related infection | 6 (32) | 2 (14) | 1 (8) | 2 (29) | |
| Soft tissue infection | 2 (11) | 2 (14) | 6 (50) | 2 (29) | |
| Osteomyelitis without implant | 2 (11) | 2 (14) | 2 (17) | | 1 (10) |
| Native joint arthritis | 3 (16) | | 1 (8) | | 2 (20) |
| Infection type, n (%) | | | | | |
| Acute | 11 (58) | 7 (50) | 8 (67) | 5 (71) | 6 (60) |
| Chronic | 8 (42) | 7 (50) | 4 (33) | 2 (29) | 4 (40) |
| Antibiotic administration prior to surgery, n (%) | | | | | |
| None | 8 (42) | 14 (100) | 4 (33) | 3 (43) | 3 (30) |
| Amoxicillin/Clavulanate | 10 (53) | | 8 (67) | 4 (57) | 5 (50) |
| Flucloxacillin | 1 (5) | | 1 (8) | | |
| Cefazolin | 3 (16) | | | 2 (29) | 4 (40) |
| Piperacillin-Tazobactam | | | | 1 (14) | 1 (10) |
| Vancomycin | 1 (5) | | 2 (17) | 1 (14) | |
| Ceftriaxon | | | 1 (8) | | |
| Ciprofloxacin | | | | | |
| Rifampicin | | | | | |
| Clindamycin | | | 1 (8) | | |
| Cefepime | | | | | 1 (10) |
| Daptomycin | | | | | 1 (10) |
| Bacteremia, n (%) | 6 (32) | 2 (14) | 1 (8) | 2 (29) | 5 (50) |
| Microbiological failure, n (%) | 6 (32) | 8 (57) | 5 (42) | 2 (29) | 7 (70) |
| Infection relapse at site | 5 (26) | 8 (57) | 5 (42) | 1 (14) | 6 (60) |
| Any previous SA infection, n (%) | 6 (32) | 2 (14) | 3 (25) | 3 (43) | 6 (60) |

C2 General
