## Supplementary figures and images for "Microgeography of staphyloccoci in human tissue explains antibiotic failure"

### Supplementary Figure 1

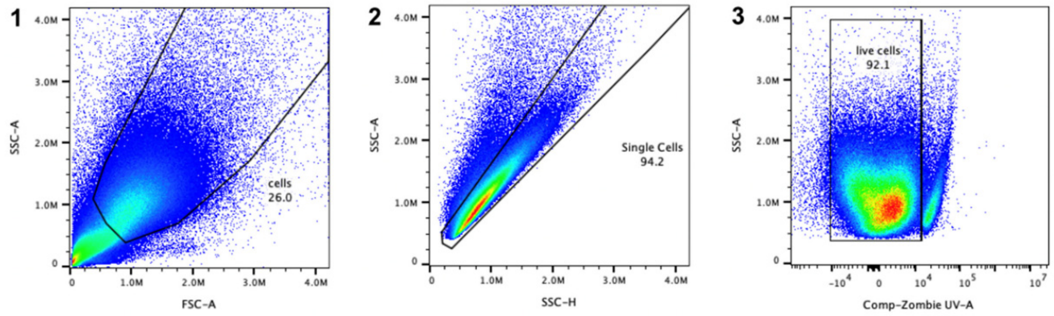


**Supplementary Figure 1: Gating strategy for flow cytometry**.
